# Sampling artifacts in single-cell genomics cohort studies

**DOI:** 10.1101/2020.01.15.897066

**Authors:** Ramon Massoni-Badosa, Giovanni Iacono, Catia Moutinho, Marta Kulis, Núria Palau, Domenica Marchese, Javier Rodríguez-Ubreva, Esteban Ballestar, Gustavo Rodriguez-Esteban, Sara Marsal, Marta Aymerich, Dolors Colomer, Elias Campo, Antonio Julià, José Ignacio Martín-Subero, Holger Heyn

## Abstract

Robust protocols and automation now enable large-scale single-cell RNA and ATAC sequencing experiments and their application on biobank and clinical cohorts. However, technical biases introduced during sample acquisition can hinder solid, reproducible results and a systematic benchmarking is required before entering large-scale data production. Here, we report the existence and extent of gene expression and chromatin accessibility artifacts introduced during sampling and identify experimental and computational solutions for their prevention.

## Introduction

Blood cells are an attractive source to systematically identify disease mechanisms and biomarkers, due to its availability in biobanks and large clinical collections. However, despite blood samples are generally archived with standardized procedures, upfront sample processing can vary profoundly even within cohorts^1^. In particular, the time between sample extraction and cryopreservation, ranging from hours (local) to days (central)^2^, might distort gene expression and epigenetic profiles and could lead to false or biased reporting. Although we have previously demonstrated that cryopreservation is a viable option for single-cell studies^3^, the effect of the sampling time on single-cell RNA (scRNA-seq) and ATAC (scATAC-seq) sequencing datasets has not been addressed.

In this work, we designed benchmarking experiments to systematically test the effect of varying processing times on single-cell transcriptome and epigenome profiles from healthy and diseased donors, while controlling for technical variability (**Online Methods**). We isolated peripheral blood mononuclear cells from healthy donors (PBMCs) and from patients affected with chronic lymphocytic leukemia (CLL), the most common adult leukemia in the Western world^4^. Samples were either preserved immediately (0 hours) or after 2, 4, 6, 8, 24 and 48 hours; simulating common scenarios in biobank and clinical routines. Single-cell 3’-transcript counting, full-length transcriptome and scATAC-seq were performed to monitor gene expression, RNA integrity and open chromatin variance across preservation time points.

## Results

### Prolonged sampling time alters single-cell transcriptome and epigenome profiles

We generated transcriptome and epigenome profiles for 66,136 and 76,146 high-quality cells, respectively. To evaluate the effect of sampling time on single-cell gene expression profiles, we initially obtained fresh PBMC from 2 healthy donors and 3 CLL patients. To simulate local processing, we stored cells prior to cryopreservation at room temperature (RT) for various time intervals up to 8h. Additionally, we stored cells for 24h and 48h, common sampling times for central sample processing. Following scRNA-seq, we detected a striking effect of the sampling time on single-cell transcriptome profiles, initiating after 2 hours and increasing in a time-dependent manner (**Fig. 1a**). This effect was reproducible across all blood cell subtypes from healthy donors and neoplastic cells from CLL patient samples (**Fig. 1a**,**b, Supplementary Fig. 1a**), and across scRNA-seq technologies (**Supplementary Fig. 1b)**. Sampling time was the major source of variability, correlating with the first principal component (PC1) for all cell subtypes (**Fig. 1c, Supplementary Fig. 1c**). Notably, although gene expression profiles varied profoundly, viable cells did not show signs of reduced RNA integrity across the time points (**Supplementary Fig. 2**).

**Figure 1.**
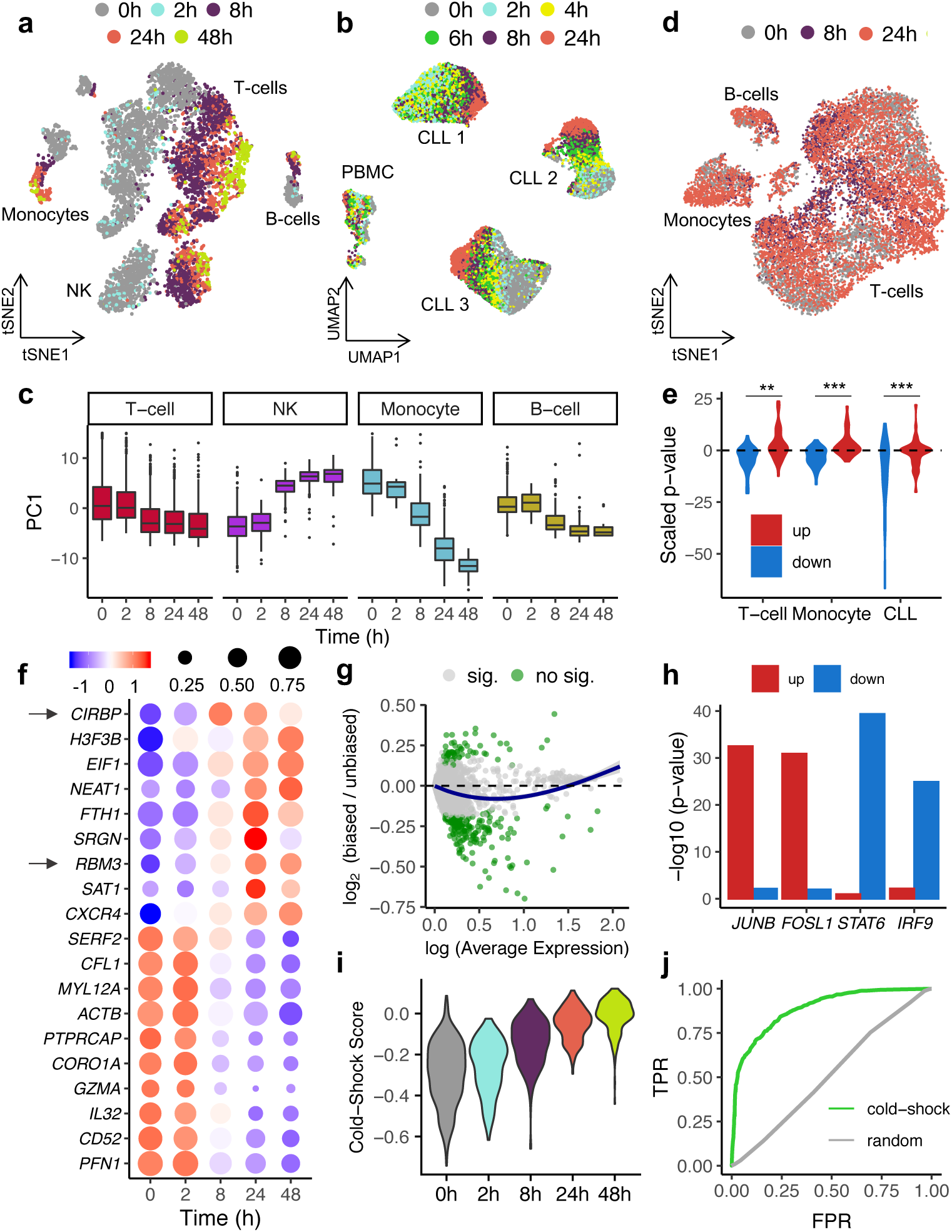
The impact of sampling time on single-cell transcriptional and open chromatin profiles. **(a, b)** scRNA-seq-based tSNE or UMAP embeddings of 7,378 PBMC (**a**, male donor) and 22,443 CLL cells (**b**, 3 donors) colored-coded by sampling time. **(c)** Distribution of the first principal component (PC1) across processing times for each PBMC subtype. **(d)** scATAC-seq-based UMAP embedding color-coded by sampling time and highlighting major PBMC cell types. (e) Violin plot showing changes in RNA expression for the 50 genes associated with the top 50 distal (enhancer) peaks changing in accessibility (down: closing sites; up: opening sites); p-value in Z-score scale, *p<0.05, **p<0.01, ***p<0.001). **(f)** Dot plot representing the time-dependent expression changes of the top up- and down-regulated genes with a minimum log(expression) of 0.5, a minimum absolute log fold-change of 0.2 and an adjusted p-value <0.001. The arrows highlight the *cold-inducible response binding protein* (*CIRBP*) and the *RNA Binding Motif Protein 3* (RBM3) genes. **(g)** M(log ratio)-A(mean average) plot showing the log2 fold-change between biased (>2h) and unbiased (<=2h) PBMC as a function of the log average expression (*Scran* normalized expression values). Significant genes are colored in green (adjusted p-value <0.001). **(h)** Motif enrichment analysis performed over the DNA sequences of the top 50 distal peaks with a change in accessibility (same peaks as **e**). **(i)** Cold-shock score distribution across processing times (female donor) calculated with the cold-shock signature defined in the male PBMC donor. **(j)** Receiver operating characteristic (ROC) curve displaying the performance of a logistic regression model in classifying “biased” and “unbiased” PBMC.

Contrary to gene expression, prolonged storage at room temperature did not cause global effects on open chromatin profiles that could be consistently detected across healthy and CLL samples (**Fig. 1d, Supplementary Fig. 3**). However, integrative analysis of scRNA-seq and scATAC-seq data pointed to a deregulation of specific genes through concerted changes at open chromatin sites. Specifically, we detected reduced expression for genes that lose open chromatin sites, a trend that was more pronounced analyzing enhancers compared to promoter sites (**Fig. 1e, Supplementary Fig. 4**) and suggesting that specific enhancer sets trigger the response to temperature changes.

### Sampling time induced a cold-shock signature in PBMC and CLL samples

Next, we aimed to determine the gene signature associated with sampling time interval to characterize, predict and correct the bias. Therefore, we conducted a differential expression analysis between affected (>2h) and unaffected conditions (≤2h). We detected 236 differentially expressed genes for PBMC (DEG, 61 up- and 175 down-regulated; **Fig. 1f**,**g, Supplementary Fig. 5a** and **Supplementary Table 1**) and 200 for CLL samples (85 up- and 115 down-regulated; **Supplementary Fig. 6a**). In addition, we observed a time-dependent decrease in the number of detected genes in both datasets (**Supplementary Fig. 6b**; p<0.001) and a global downregulation of gene expression (**Fig. 1g, Supplementary Fig. 6a**). This global effect has been reported previously in bulk transcriptomics studies^5^, pointing to a reduction of the transcriptional rate when cells are removed from their physiological niche (37°C) and stored at RT (21°C).

In line with the above findings, the sampling time-associated transcriptional signature exhibited three defining characteristics of a cold-shock response. First, the *Cold Inducible RNA Binding Protein* (*CIRBP*) and the *RNA Binding Motif Protein 3* (*RBM3*), two cold-shock master regulators^6^, were up-regulated in a time-dependent manner (**Fig. 1f, Supplementary Fig. 6a**). Second, genes involved in a negative regulation of translation (including *EIF1* and *EIF1B*) were induced upon cold exposure (**Fig. 1f, Supplementary Fig. 6a**,**c**), in agreement with the blockade of 5’cap-dependent translation under such conditions^7^. Finally, down-regulated genes were enriched in “Arp2/3 complex-mediated actin nucleation” processes, supporting a cold-shock-dependent depolymerization of the cytoskeleton (**Supplementary Fig. 6c**)^8^. In addition to the cold-shock response, we observed a pronounced down-regulation of immune cell type specific genes (**Fig. 1f, Supplementary Fig. 6a**) and programs (**Supplementary Fig. 5b**,**c** and **6d**) pointing to a loss of identity in prolonged storage conditions. Motif enrichment analysis at sampling time-sensitive enhancers identified by scATAC-seq pointed to a significant increase in the accessibility of transcription factor binding sites (TFBS) of early stress response genes, such as *JUNB* and *FOSL1* (**Fig. 1h, Supplementary Table 1**), as previously shown in scRNA-seq studies^9^. Further, we detected a significant decrease in accessibility at TFBS of immune and inflammation-related genes, such as *STAT6* and *IRF9* (**Fig. 1h, Supplementary Table 1**), in line with the downregulation of immune response genes at the transcript level. Finally, to predict such sampling time-effect, we calculated a cold-shock score using the abovementioned signature^10^, which classified cells to be affected by sampling time (AUC = 0.888, **Fig. 1i**,**j**).

### Correcting and preventing sampling biases

We sought to identify solutions for retrospective study designs and prospective cohort collection. *In silico* data correction is commonly applied to diminish the effects of technical or biological variability in scRNA-seq datasets by scoring and regressing out specific gene sets^11^. Applying such strategy on the cold-shock gene expression score, we were able to reduce the sampling effect, especially for samples with local processing (≤8h). This correction was robust for different PBMC subtypes (Kbet score^12^; **Fig. 2a**,**b**) and neoplastic cells from CLL patients (**Supplementary Fig. 7a**) as well as simulated datasets with varying proportions of affected cells (**Supplementary Fig. 7b**), suggesting a broad application spectrum. Importantly, the correction conserved biological variance related to cell identity in blood and inter-individual variation in CLL patients. However, owing to the Simpson’s paradox^13^ and to gene expression pleiotropy^14^, regressing out technical confounders can remove subtle biological heterogeneity and homogenize cell subpopulations, which can challenge data interpretation. Consequently, we sought experimental alternatives to reduce sampling effects in retrospective study designs. We reasoned that the magnitude of gene expression alterations can be diminished by cell culture and through the activation of cell type specific programs. Hence, we utilized PBMCs (cryopreserved at 0/8/24 hours) and processed them directly (day 0) or after two days in cell culture with simultaneous T-cell activation (anti-CD3, day 2). Strikingly, the culturing/activation reduced the sampling induced artifact, quantifiable through increased similarities between the time points (Kbet score^12^; **Fig. 2c**,**d**). In line, after cell culture, no significant differences in cold-shock score could be observed between the time points (**Supplementary Fig. 8**).

**Figure 2.**
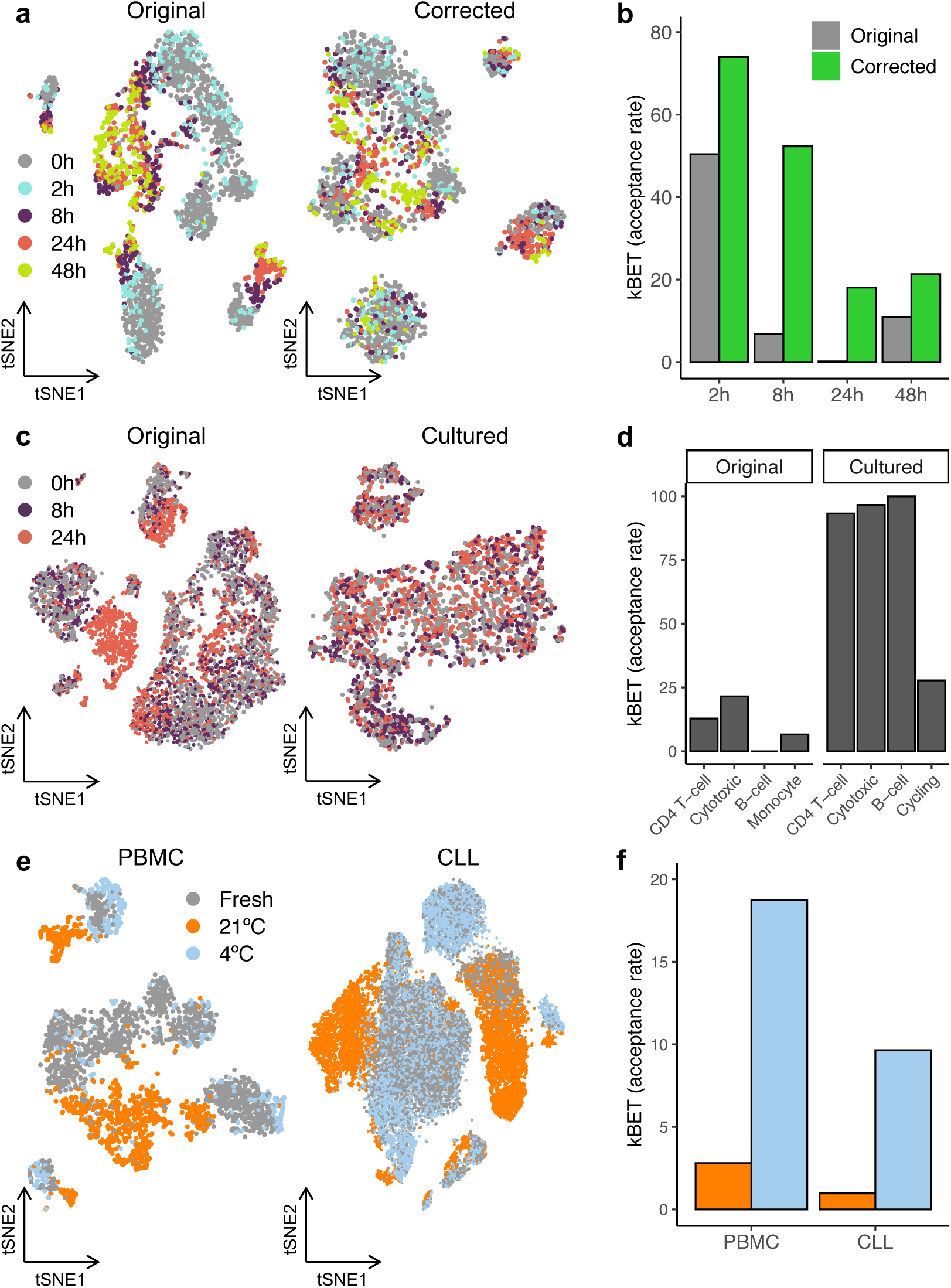
Solutions to correct or prevent sampling time-induced artifacts. **(a)** tSNEs displaying the effect of varying processing times on the transcriptome profiles of 7,378 PBMC before (left) and after (right) regressing out the cold-shock score for every highly variable gene. **(b)** kBET acceptance score distribution across sampling times with or without the computational correction. **(c)** tSNE showing the effect of PBMC culturing and activation with anti-CD3 Dynabeads over two days. **(d)** kBET acceptance score distribution across cell types with or without cell cultur/activation. **(e)** tSNE highlighting the sampling effect between cells cryopreserved immediately (fresh, 0 h) or after 24 h and 48 h stored cold (4°C) or at RT (21°C). **(f)** kBET acceptance score distribution across storage temperatures.

Finally, we hypothesized that cold sample storage could prevent time-related sampling effects by minimizing active and passive cell responses. In line, tissues (lung, pancreas and oesophagus) preserved at cold temperatures (4°C) did not show altered single-cell gene expression profiles or cell type composition changes up to 72 hours of storage^15^. Importantly, changing storage temperatures could be readily implemented in prospective cohort study designs to enable subsequent scRNA-seq. Indeed, when PBMC or CLL samples were stored at 4°C until cryopreservation (24/48 h), we did not detect global gene expression artifacts; an effect observed for both healthy and CLL samples and replicated across donors and technologies (**Fig. 2e**,**f, Supplementary Fig. 9**).

## Conclusions

We report that varying sampling times until cryopreservation is a driver of technical variability in scRNA-seq and scATAC-seq profiles. This bias was ubiquitous across cell types, donors, protocols and disease status, thus, likely presents a highly frequent obstacle in transcriptome and epigenome cohort studies. Despite the substantial impact on single-cell datasets, computational corrections, cell culture and storage adjustments allow the design of reliable retro- or prospective studies. The here detected artifacts are important to consider when planning single-cell experiments. Failing to select suitable samples or to correct datasets will lead to biased or false reporting.

## Acknowledgements

HH is a Miguel Servet (CP14/00229) researcher funded by the Spanish Institute of Health Carlos III (ISCIII). CM and MK are supported by AECC postdoctoral fellowships. This work has received funding from the Ministerio de Ciencia, Innovación y Universidades (SAF2017-89109-P; AEI/FEDER, UE). This study was further funded by the Spanish Ministry of Economy and Competitiveness (grant number: IPT-010000-2010-36, cofunded by the European Regional Development Fund). Core funding is from the ISCIII and the Generalitat de Catalunya. We acknowledge support of the Spanish Ministry of Economy, Industry and Competitiveness (MEIC) to the EMBL partnership, the Centro de Excelencia Severo Ochoa, the CERCA Programme / Generalitat de Catalunya, the Spanish Ministry of Economy, Industry and Competitiveness (MEIC) through the Instituto de Salud Carlos III and the Generalitat de Catalunya through Departament de Salut and Departament d’Empresa i Coneixement. We also acknowledge the Co-financing by the Spanish Ministry of Economy, Industry and Competitiveness (MEIC) with funds from the European Regional Development Fund (ERDF) corresponding to the 2014-2020 Smart Growth Operating Program. We acknowledge the Generalitat de Catalunya Suport Grups de Recerca AGAUR 2017-SGR-736 (to JIMS) and 2017-SGR-1142 (to EC), and CIBERONC (CB16/12/00225 and CB16/12/00334). EC is an ICREA Academia Researcher. This project received support from the European Commission under the projects DocTIS (H2020, SEP-210574908). This publication is part of a project (BCLLATLAS) that has received funding from the European Research Council (ERC) under the European Union’s Horizon 2020 research and innovation programme (grant agreement No 810287).

## Ethical Statement

The enrolled CLL patients gave informed consent for this study following the ICGC guidelines and the ICGC Ethics and Policy committee. This study was approved by the clinical research ethics committee of the Hospital Clínic of Barcelona. For healthy PBMC donors, the study was approved by the local ethics committee and all the patients signed an informed consent. This study was conducted in accordance with the Declaration of Helsinki principles.

## Author’s Contributions

HH, IMS and AJ designed the study. RMB performed all gene expression data analyses. GI performed the ATAC-seq analysis. CM, DM, MK, JRU and NP prepared the PBMC and CLL samples. GRE supported the data analysis. SM, AJ, EC and IMS provided samples and additional support. HH and RMB wrote the manuscript. All authors read and approved the final version.

## Code and Data Availability

The analysis notebooks and feature-barcode matrices to reproduce the aforementioned analysis are hosted at https://github.com/massonix/sampling_artifacts. The complete raw data will be available at the Human Cell Atlas Data Coordination Portal (DCP).

## Online Methods

### PBMC isolation and cryopreservation

Peripheral venous blood samples were collected by venipuncture from two voluntary blood donors, one male and one female. Blood samples were collected in ACD-tubes and stored at room temperature (RT) or 4°C. In the former condition peripheral blood mononuclear cells (PBMCs) were isolated at 2, 8, 24 and 48 h. For samples stored at 4°C, PBMCs were isolated at 24 and 48hrs. PBMCs separation was performed using Ficoll density gradient centrifugation. For each condition, 12 ml of blood were diluted with an equal volume of pre-warmed RPMI 1640 culture medium (Lonza). The diluted blood was then carefully layered onto a Leucosep tube (Greiner Bio-One) prefilled with 15 ml of Ficoll-Plus (GE Healthcare Biosciences AB) and centrifuged for 15 min at 800 x g and RT (without acceleration and brake). After centrifugation, PBMCs were collected with a sterile Pasteur pipette into a 50 ml tube, diluted up to 10 ml with pre-warmed RPMI medium and centrifuged for 10 min, at 400 x g and RT. Following a second washing step with 5 ml of RPMI medium and a 5 min centrifugation, PBMCs were resuspended in 8 aliquots of freezing media. Freezing media consisted of RPMI 1640 with 20% heat-inactivated fetal bovine serum (Sigma-Aldrich), 10% DMSO (Sigma-Aldrich) and Penicillin-Streptomycin 1:1000 (Lonza). 1 ml aliquots, with approximately 1×10e6 cells/ml, were gradually frozen using a commercial freezing box (Mr. Frosty, Nalgene) at −80°C for 24 h and then stored in a vapor– phase liquid nitrogen tank at −150°C.

Cryopreserved (−80°C) PBMC samples were rapidly thawed in a 37°C water bath. Each sample was transferred into a 15 ml Falcon using a 1000 ul cut tip without mixing by pipetting. Next, 1 ml of 37°C pre-warmed media (Hibernate-A supplemented with 10% FCS; ThermoFisher) was added drop-wise with gently swirling of the sample. After 1 min incubation, 2 ml of pre-warmed media were added and incubated for 1 min. Next, 5 ml pre-warmed media was gently added, inverted and incubated (1 min). This step was repeated once. Finally, the samples were centrifuged at 700 x g for 5 min (4°C). The supernatant was removed and the pellets re-suspended in 100 ul of Cell staining buffer (BioLegend). CLL patient samples were obtained from freshly extracted blood, stored either at RT or 4°C. Mononuclear cells were isolated after Ficoll density gradient centrifugation, at 2, 4, 6, 8 and 24 h after patient blood extraction. The cells were directly cryopreserved with freezing media (RPMI 1640 with 20% FBS and 10% DMSO), in the concentration of 5-10×10e6 cells/ml, according to standarized protocol. The tumor cell content of all the samples was >80%, as assassed by immunostaining of CD19, CD20, CD5 and CD45 followed by flow cytometry. All patients gave informed consent for their participation in the study according to International Cancer Genome Consortium (ICGC) guidelines. Cell hashing was performed following manufacturer’s instructions (Cell hashing and Single Cell Proteogenomics Protocol Using TotalSeq™ Antibodies; BioLegend). Therefore, samples were incubated 10 min at 4°C with Human TruStain FcX™ Fc Blocking reagent (BioLegend). Next, sample-specific TotalSeq antibodies (anti-human Hashtag 1-8, Biolegend) were added with subsequent incubation on ice for 45 min. Cells were washed once with cold 1X PBS supplemented with 0.0005% BSA (ThermoFisher) and pelleted at 700 x g for 5 minute. A single cell solution was obtained resuspending the pellet in 1X PBS (0.0005% BSA) and filtering it through a 40 µm cell strainer. The cells were counted in an automatic cell counter (Countess® v.2, ThermoFisher).

### PBMC isolation and activation

Peripheral venous blood samples were collected by venipuncture from three voluntary donor (1 male and 2 females). Blood samples were collected in 10 ml Vacutech Vacuum Blood Collection Tubes K2/K3 EDTA (Becton Dickinson) and stored at RT. Peripheral blood mononuclear cells (PBMCs) were isolated at 0, 8 and 24 hours after blood collection. PBMCs separation was performed using Ficoll density gradient centrifugation. For each condition, 9 ml of blood were diluted with an equal volume of 1X PBS (Gibco). The diluted blood was then carefully layered onto 9 ml of Lymphoprep solution (STEMCELL Technology) and centrifuged for 15 min at 700 x g and RT (without acceleration and brake). After centrifugation, PBMCs were collected and washed twice with 10 ml of 1X PBS. The pellet was resuspended with 10 ml of 1X PBS and cells were counted with a TC20™ Automated Cell Counter (Bio-Rad Laboratories). PBMCs were again centrifuged for 5 min at 700 x g and resuspended in an appropriate volume of freezing media (RPMI with 10% heat-inactivated fetal bovine serum and 10% DMSO). Aliquots of ∼0.5 x 10e6 cells/ml were gradually frozen using a commercial freezing box (Mr. Frosty, Nalgene) at −80°C for 24 h and then stored in a vapor–phase liquid nitrogen tank at −150°C. For T-cell activation, cells were thawed in MACS buffer (1X PBS, 4% FBS, 2 mM EDTA), centrifuged during 5 min at 700 x g and RT, and resuspended in pre-warmed culture media (RPMI, 1% Pyruvate, 20% FBS, Pen/Strep, DNase 100 U/ml). A TC20™ automated cell counter was used to assess cell number and viability. The number of only viable cells was used to calculate volumes for cell seeding. For each condition, 200,000 live cells were seeded into two wells of a 96-well round bottom plate (Sigma Aldrich) for a total of 400,000 cells per condition (time point). Dynabeads Human T-Activator CD3/CD28 (Thermo Fisher Scientific) were transferred to a 1.5 ml tube (5 ul/well), washed twice with 1 ml of cell culture media and resuspended with 10 volumes of cell culture media. 50 ul of resuspended beads were added to each well for T-cell activation and expansion. Cells were incubated during 24 hours at 37°C with 5% CO2 and 5% humidity. The remaining cells (∼350,000 cells per condition) were used as a control (day 0) for T-cell activation. Cells subjected to T-cell activation protocol were collected in a 1.5 ml tube and stained with DAPI (Thermo Fisher Scientific) at 1 µM final concentration. DAPI-negative live individual cells were sorted with a BD FACSAria™ Fusion Flow cytometer (BD Biosciences) in 1X PBS supplemented with 0.05% BSA.

Samples subjected to T-cell activation treatment, as well as corresponding control samples, were subjected to a Cell Hashing protocol before proceeding to scRNA-seq. Cell hashing was performed following manufacturer’s instructions (Cell hashing and Single Cell Proteogenomics Protocol Using TotalSeq™ Antibodies; BioLegend). Cells were counted with a TC20™ Automated Cell Counter, and an equal number of cells was taken for each condition. Briefly, samples were resuspended in Cell Staining Buffer (BioLegend), incubated 10 min at 4°C with Human TruStain FcX™ Fc Blocking reagent (Bio Legend). To each condition, a specific TotalSeq-A antibody-oligo conjugate (anti-human Hashtag 1-8, Biolegend) was added and incubated on ice for 1 hour. Cells were then washed three times with cold PBS-0.05% BSA (ThermoFisher) and centrifuged for 5 min at 700 x g at 4°C. Finally, cells were resuspended in an appropriate volume of 1X PBS-0.05% BSA in order to obtain a final cell concentration >500 cells/ul, suitable for 10x Genomics scRNA-seq. An equal volume of hashed cell suspension from each of the conditions (0 h, 8 h and 24 h) was mixed and filtered with a 40 µm strainer. Cell concentration was verified by counting with a TC20™ Automated Cell Counter.

### Single-cell RNA sequencing

Cells were partitioned into Gel Bead In Emulsions with a Target Cell Recovery of 10,000 total cells. Sequencing libraries were prepared using the single-cell 3’ mRNA kit (V2 and V3; 10X Genomics) with some adaptations for cell hashing, as indicated in TotalSeq™-A Antibodies and Cell Hashing with 10x Single Cell 3’ Reagent Kit v3 3.1 Protocol by BioLegend. Briefly, 1 µl of 0.2 µM HTO primer (Hashtag Oligonucleotides; GTGACTGGAGTTCAGACGTGTGC*T*C; *Phosphorothioate bond) was added to the cDNA amplification reaction in order to amplify the hashtag oligos together with the full-length cDNAs. A SPRI selection clean-up was done in order to separate mRNA-derived cDNA (>300 bp) from antibody-oligo-derived cDNA (<180 bp), as described in the above mentioned protocol. 10x cDNA sequencing libraries were prepared following 10x Genomics Single Cell 3’ mRNA kit protocol, while HTO cDNAs were indexed by PCR as follows. Briefly, 5 µl of purified hashtag oligo cDNA were mixed with 2.5 µl of 10 µM Illumina TruSeq D70X_s primer (IDT) carrying a different i7 index for each sample, 2.5 µl of SI primer from 10x Genomics Single Cell 3’ mRNA kit, 50 µl of 2X KAPA HiFi PCR Master Mix (KAPA Biosystem) and 40 µl of nuclease-free water. The reaction was carried out using the following thermal cycling conditions: 98°C for 2 min (initial denaturation), 12 cycles of 98°C for 20 sec, 64°C for 30 sec, 72°C for 20 sec, and a final extension at 72°C for 5 min. The HTO libraries were purified with 1.2 X SPRI bead selection. Size distribution and concentration of cDNA and HTO libraries were verified on an Agilent Bioanalyzer High Sensitivity chip (Agilent Technologies). Finally, sequencing of HTO and cDNA libraries was carried out on a HiSeq4000 or NovaSeq6000 system (Illumina).

### Single-cell ATAC sequencing

For the single-cell ATAC-seq experiments, we analyzed one PBMC and one CLL sample isolated after 0 h, 8 h and 24 h of blood storage at room temperature before cryopreservation. Frozen samples were rapidly thawed in a 37°C water bath. Each sample was transferred into a 15 ml Falcon using a 1000 µl cut tip without mixing by pipetting. Next, 1 ml of 37°C pre-warmed media (Hibernate-A supplemented with 10% FCS) was added drop-wise with gentle swirling of the sample. After 1 min of RT incubation, additional 2 ml of pre-warmed media were added. The samples were again kept at RT for 1 min, before 5 ml of pre-warmed media were gently added. This step was repeated once. Then, samples were centrifuged at 700 x g for 5 min. Supernatant was removed and the pellets resuspended in 500 µl of PBS supplemented with 0.05% BSA. Cell concentration and viability were determined with a TC20™ Automated Cell Counter.

Nuclei isolation was performed following the “Nuclei Isolation for Single Cell ATAC Sequencing demonstrated protocol” (10x Genomics). Briefly, 1,000,000 cells from the CLL sample and 300,000 cells from PBMCs, were transferred to a 1.5 ml microcentrifuge tube and centrifuged at 500 x g for 5 min at 4°C. The supernatant was removed without disrupting the cell pellet and 100 µl of chilled Lysis Buffer (10 mM Tris-HCl (pH 7.4); 10 mM NaCl; 3 MgCl2; 0.1% Tween-20; 0.1% Nonidet P40 Substitute; 0.01% Digitonin and 1% BSA) were added and pipette-mixed 10 times. Samples were then incubated on ice during 3 min. Following lysis, 1 mL of chilled Wash Buffer (10 mM Tris-HCl (pH 7.4); 10 mM NaCl; 3 MgCl2; 0.1% Tween-20 and 1% BSA) was added and pipette-mixed. Nuclei were centrifuged at 500 x g for 5 min at 4°C and the supernatant removed without disrupting the pellet. Based on the starting number of cells and assuming a 50% loss during the procedure, nuclei were resuspended into the appropriate volume of chilled Diluted Nuclei Buffer (10x Genomics) in order to achieve a nuclei concentration of 1,540-3,850 nuclei/µl, suitable for a Target Nuclei Recovery of 5000. The resulting nuclei concentration was determined using a with a TC20™ Automated Cell Counter.

scATAC-seq libraries were prepared according to the Chromium Single Cell ATAC Reagent Kits User Guide (10x Genomics; CG000168 Rev B). Briefly, the transposition reaction was prepared by mixing the desired number of nuclei with ATAC Buffer (10X Genomics) and ATAC Enzyme (10X Genomics), before incubation for 60 min at 37°C. Nuclei were partitioned into Gel Bead-In-Emulsions (GEMs) by loading the following into a Chip E: the master mix (previously added to the same tube of the transposed nuclei), the Chromium Single Cell ATAC Gel Beads (10X Genomics) and the Partitioning Oil (10X Genomics). After the run into the Chromium Controller, the DNA linear amplification was performed by incubating the GEMs at the following thermal cycling conditions: 72°C for 5 min, 98°C for 30 sec, 12 cycles of 98°C for 10 sec, 59°C for 30 sec and 72°C for 1 min. GEMs were broken using the Recovery Agent (10X Genomics), and the resulting DNA was purified by Dynabeads and SPRIselect reagent (Beckman Coulter; B23318) bead clean-ups. Indexed sequencing libraries were obtained by mixing the amplification product with the Sample Index PCR Mix (10X Genomics) and the Chromium i7 Sample Index (10x Genomics), and incubating at the following thermal cycling conditions: 98°C for 45 sec, 12 cycles of 98°C for 20 sec, 67°C for 30 sec, 72°C for 20 sec with a final extension of 72°C for 1 min. Sequencing libraries were subjected to a final bead clean-up SPRIselect reagent and quantified on an Agilent Bioanalyzer High Sensitivity chip (Agilent Technologies). Finally, libraries were loaded on an Illumina HiSeq 2500 system in Rapid Run mode using the following read length: 50 bp Read 1N, 8 bp i7 Index, 16 bp i5 Index and 50 bp Read 2N.

### Primary processing and demultiplexing

We processed sequencing reads with CellRanger v3.0.0 for the PBMC data and v3.0.2 for the CLL and T-cell activation data. We used the human GRCh38 assembly as reference genome. To specify the hashtag oligonucleotide (HTO) libraries, the cDNA libraries and the HTO sequences, we followed the “Feature Barcoding Analysis” pipeline, available at https://support.10xgenomics.com/single-cell-gene-expression/software/pipelines/latest/using/feature-bc-analysis. We set the --chemistry and --expect-cells flags of *cellranger count* to “SC3Pv3” and “5000”, respectively. We demultiplexed cell hashtags as described in Stoeckius et al.^16^ for each batch and donor separately. Briefly, we normalized hashtag oligonucleotide (HTO) counts using a centered log ratio (CLR), in which each count is divided by the geometric mean of a HTO across cells and log-transformed. We then clustered barcodes using k-medoids with k equal to the number of conditions (k=4 for batch 03, and k=8 for batch 04), which allowed us to identify the background distribution of each HTO. We re-clustered Male 04 with k=3, as no clear signal for the HTO “24h 4°C” was detected. Subsequently, we considered the top 0.5% normalized HTO counts of the background distribution as outliers and excluded them. We classified barcodes to a given condition if the normalized HTO counts of that condition exceeded the 0.99 quantile. We discarded both barcodes that were assigned to more than one condition (multiplets) and barcodes that were not assigned to any condition (negatives). In subsequent datasets (CLL and T-cell activation), we demultiplexed the HTO with Seurat’s built-in functions^17^.

### Quality control and normalization

We performed quality control (QC) and normalization separately for each dataset (PBMC, CLL, T-cell activation). Following the guidelines from Luecken et al.^18^, we inspected the distributions of three main QC metrics: library size (total UMI), library complexity (number of detected genes) and mitochondrial expression. Importantly, we analyzed these metrics jointly to ensure that cells with high mitochondrial expression were not metabolically active. Finally, we analyzed one of the CLL donors independently as it showed markedly different distributions in QC metrics. We classified as damaged cells those barcodes with an aberrantly low number of UMI and genes, or with an abnormally high mitochondrial expression. Likewise, we classified as doublets those barcodes that possessed and aberrantly large library size and complexity. We also leveraged DoubletFinder^19^ to detect doublets that shared the same HTO and were not outliers in any qc metric. We ruled out genes that were detected in less than 10 (CLL) or 15 cells (T-cell activation). Finally, we used the *Scran* 1.10.2 package^20^ to normalize UMI counts with cell-based size factors.

### Cell type annotation

Cell type annotation was performed within the *Seurat* framework^21^ (**Supplementary Fig. 10**). To cluster cells, we: (i) identified overdispersed genes with the *FindVariableFeatures* function (using default parameters), (ii) scaled UMI counts and regressed out the batch effect, (iii) performed Principal Component Analysis (PCA), (iv) used these PCs to create a k-nearest neighbors graph with the *FindNeighbors* function and (v) clustered cells with the *FindClusters* function. We set the resolution parameter to 0.05, 0.01 and 0.1 for the PBMC, CLL and T-cell activation data, respectively. Finally, we used well-known marker genes to annotate each cluster to its specific cell type.

### Variance analysis

To elucidate which variables introduced more variability on the expression matrix, we used the *plotExplanatoryVariables* function from *Scater*^22^, which fits a linear model for each gene with only one confounding factor (i.e. detected genes) as explanatory variable. Then, the distribution of R^2^ values (% of explained variability) for the variables with the largest R^2^ is plotted.

### Differential expression analysis (DEA)

To find the cold-shock signature, we divided cells in the PBMC and CLL datasets in time-biased (t >2h) and time-unbiased (t <=2h). Subsequently, we performed a Wilcoxon signed-rank test to test for differential expression for each gene. Soneson et al.^23^ reported that this test is among the best performing for scRNA-seq DEA analysis. Vieth et al.^24^ showed that with *Scran* normalization, there is no need for scRNA-seq-tailored statistical tests. We defined as signature those genes with an adjusted p-value <0.001.

### Gene Ontology (GO) enrichment analysis

To elucidate biological processes affected by sampling time, we conducted a GO enrichment analysis with the *GOstats* 2.48.0 package^25^. We used as target set the entrez identifiers of the up-regulated (log fold-change >0) or down-regulated (log fold-change <0) genes in the cold-shock signature, and as universe set the entrez identifiers of all genes that we included in the analysis. Finally, we filtered out GO terms that were too general (Size >=300), or too specific (Size <3). In addition, we only retained GO terms with a p-value lower than 0.05 and an odds ratio greater than two.

### Prediction of cold-shocked cells

To predict cold-shocked cells, we used the *AddModuleScore* function of Seurat to compute a cold-shock score per cell using a signature calculated on the male donor (training set). We then fitted a logistic regression model using the cold-shock score as explanatory variable. Subsequently, we predicted the probability of being “biased” for every cell of the female donor (test set), and found the Area Under the Curve (AUC) with the “AUC” function from the cvAUC v1.1.0 package. To test the significance of our signature, we repeated the process with a signature defined on random genes.

### Computational correction cold-shock signature

To correct the time-biased transcriptomes in the female PBMC dataset (test set), we regressed the expression of each gene on the cold-shock score, and kept the scaled and centered residuals as the variance in gene expression not explained by time. We performed this process for each cell type independently to minimize Simpson’s paradox.

As all the analysis above used a similar proportion of “biased” and “unbiased” cells, we sought to test the effect of varying percentages of biased cells on the cold-shock score computation and regression. In this setting, we performed bootstrapping as follows: first, we sampled 300 cells with replacement from the Smart-seq2 dataset, enforcing an approximate percentage of time-affected cells. Second, we computed the average Silhouette width between affected and unaffected cells. Finally, we computationally corrected the transcriptome profiles and recalculated the average Silhouette width. We repeated this process 25 times for each of a set of percentages ranging from 10% to 90% of affected cells.

### k-nearest-neighbor batch-effect test (kBET)

To assess the mixability between cells of different time-points in the presence or absence of our corrections, we used the kBET metric^12^. Briefly, kBET compares the label distribution of the local k-nearest neighborhood of a given cell with the global distribution with a Pearson’s χ^2^ test, with the null hypothesis that, if samples are well-mixed, both distributions are equal. We ran kBET with the cells embedded in UMAP space and with default parameters. We defined the acceptance rate as the percentage of tested cells with a p-value >0.05, and as rejection rate 100-acceptance rate.

### Smart-seq2 validation

To confirm that the results obtained from 10x Genomics-derived data were technology-independent, we profiled the transcriptome of 376 CD3+ T-cells with Smart-seq2^26^. The cells originated from the same donors as in the 10x Genomics experiments, and were distributed across four 96-well plates (all time points per plate). We discarded 60 cells that either had <75,000 or >1,000,000 total counts, <435 detected genes or a mitochondrial expression >20%. Similarly, we filtered out 6,542 genes that had an average expression across cells <1. We normalized gene counts with the *Scran* package, which removed the batch effect between plates. Finally, we clustered cells with the *SC3* 1.10.1 package^27^, as it outperforms other tools for small datasets^28^.

### ATAC-seq data analysis

ATAC-seq data from 10x Genomics was processed with CellRanger-atac 1.1.0. Differential accessibility to detect changes in open chromatin sites was performed with Wilcoxon-Mann-Whitney Rank Sum Test for High-Throughput Expression Profiling Data (R BioQC v1.0.0). Motif enrichment analysis was performed using the package *motifcounter* v.1.10.0^29^ with default parameters and the motifs were downloaded from JASPAR database (579 motifs from JASPAR CORE VERTEBRATES, http://jaspar.genereg.net/downloads/). The background distribution was computed over the total peaks called in the datasets (56,627 in the PBMCs and 80,861 in the CLL).

## Supplementary Material

**Supplementary Figure 1.**
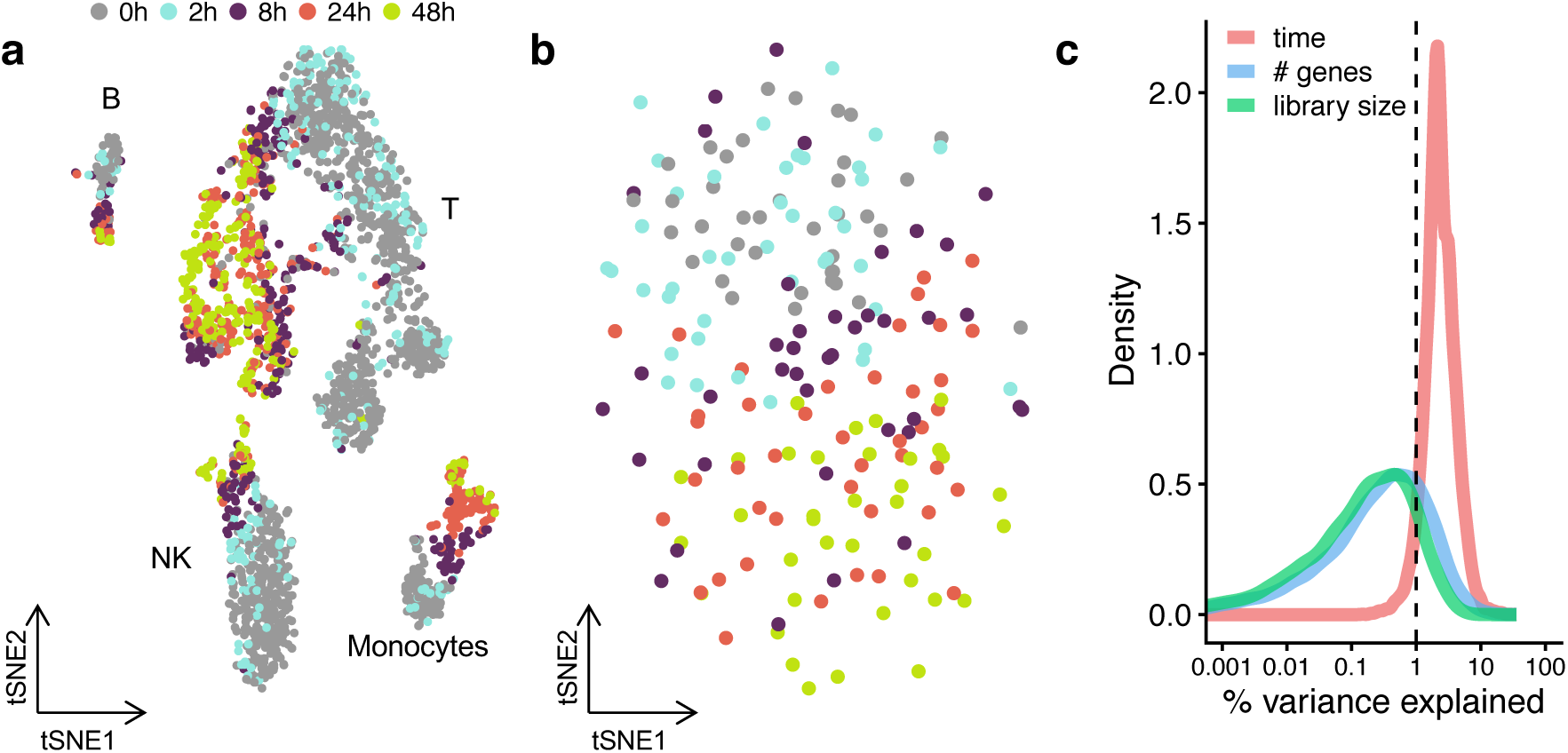
The impact of sampling time on single-cell transcriptional profiles. **(a**,**b)** tSNE embedding of **(a)** 2,460 PBMC (10x Genomics, female donor) and **(b)** 198 CD3-positive cells (Smart-seq2) colored by sampling time. (**c**) Distribution of R^2^ obtained by regressing the expression of each gene in PBMCs onto different technical variables (sampling time, library size and number of detected genes).

**Supplementary Figure 2.**
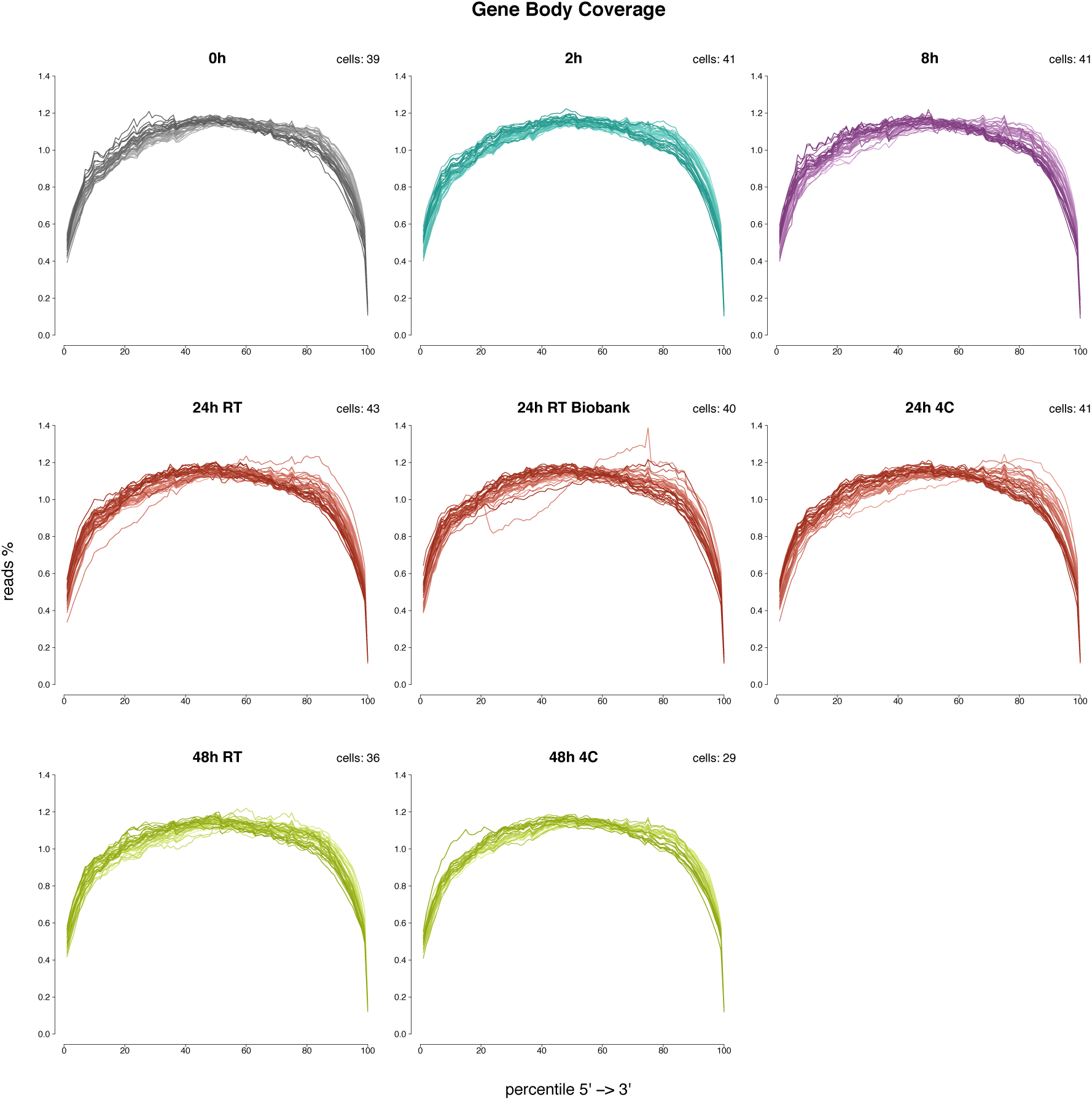
Conserved RNA integrity across sampling time points. Mapping distribution of sequencing reads from 5’ to 3’ for full-length single-cell library preparation from 198 CD3-positive cells (Smart-seq2) across different time-points. Each line represents the distribution of a single cell. The total number of cells per time point is indicated.

**Supplementary Figure 3.**
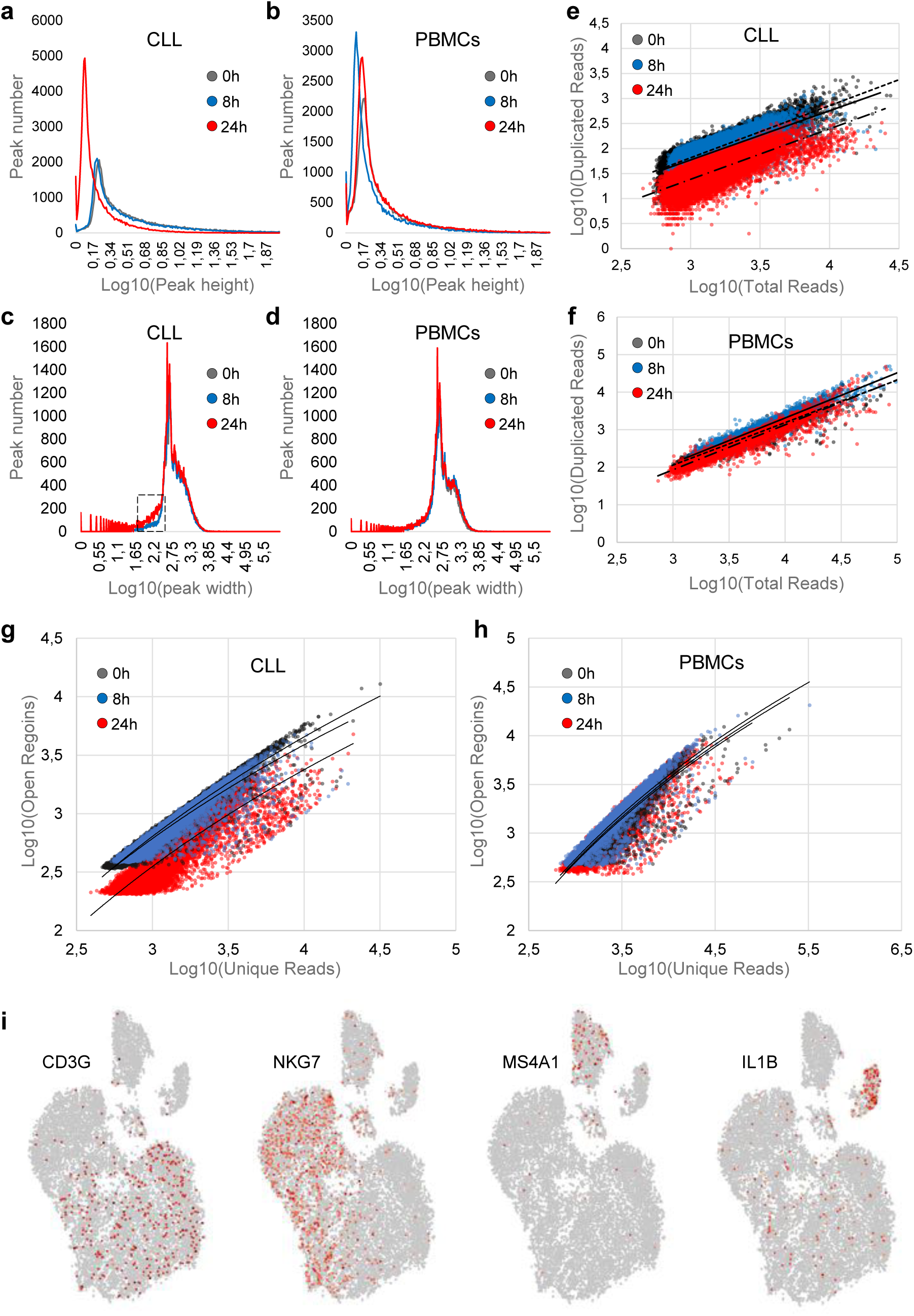
Single-cell ATAC-seq data analysis. (**a**,**b**) Distribution of peak heights in the different conditions. Peak heights were estimated by summing the reads of all the cells of a given condition and dividing by the peak width (bp). The CLL 24 h sample displays an abundance of small peaks compared to 0 and 8 hours, an effect not observed PBMC datasets, pointing to be a technical rather than sampling bias. (**c**,**d**) Distribution of peak widths in the different conditions. CLL 24 h sample had narrower peaks (visible especially in the highlighted range). (**e**,**f**) Relationship between total sequencing reads and duplicated reads at single-cell resolution. Duplicated reads originate from PCR amplification of the same amplicon. A lower amount of duplicated reads at a given sequencing depth, as observed in the CLL 24 h sample, indicates higher background signal. (**g**,**h**) Relationship between unique sequencing reads (ie, after removal of duplicate reads) and detected open regions (ie peaks) at single-cell level. The CLL 24 h sample had less detected open regions for the same amount of unique reads, again compatible with a higher background noise. (**i**) t-SNE plots of scATAC-seq dataset of PBMCs showing the expression of CD3G (T-cells), NKG7 (Natural killer cells), MS4A1 (B-cells) and IL1B (Monocytes).

**Supplementary Figure 4.**
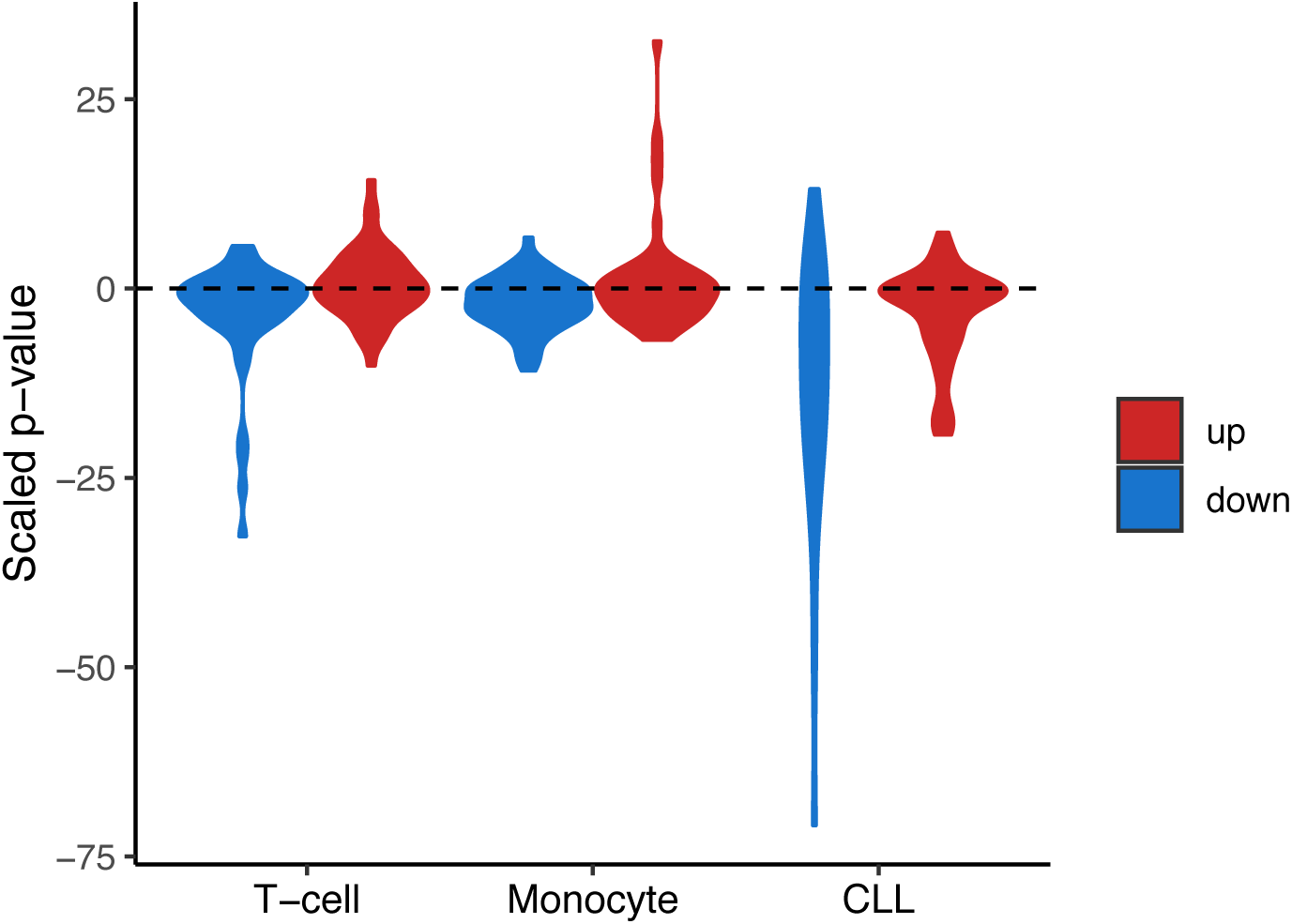
Violin plots showing the changes in RNA expression for the 50 genes associated with the top 50 promoter peaks with a change in accessibility (UP and DOWN, p-value in Z-score scale).

**Supplementary Figure 5.**
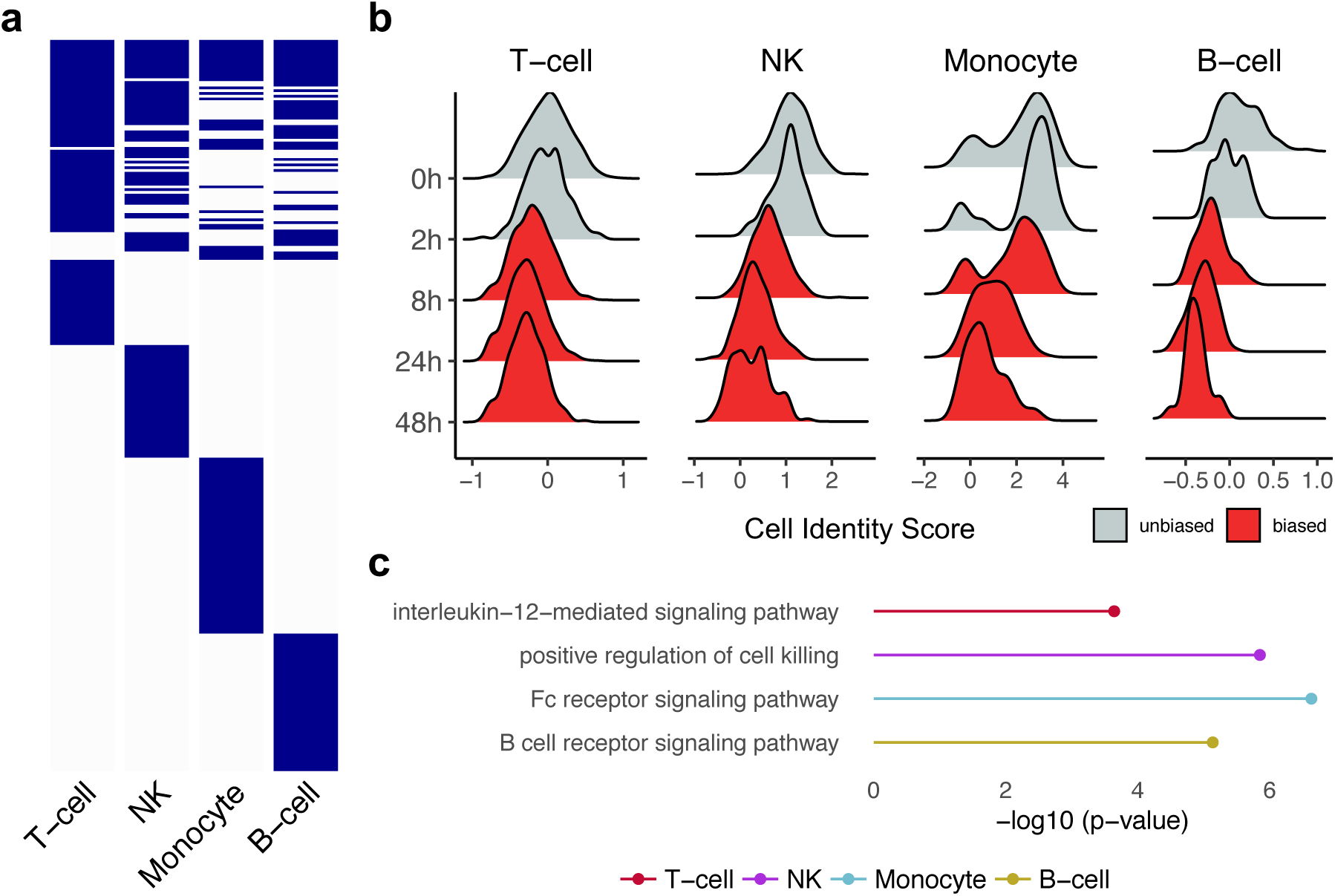
Sampling time induces a loss of cell identity in PBMC. (**a**) Heatmap showing the cell type specificity of the cold-shock transcriptome signature (top 100 differentially expressed genes per cell type). (**b**) Time-dependent loss of cell identity for each PBMC subtype. Cell identity scores are calculated using manually curated cell subtype markers. (**c**) Gene ontology enrichment analysis of the down-regulated genes in each cell type-specific cold-shock signature.

**Supplementary Figure 6.**
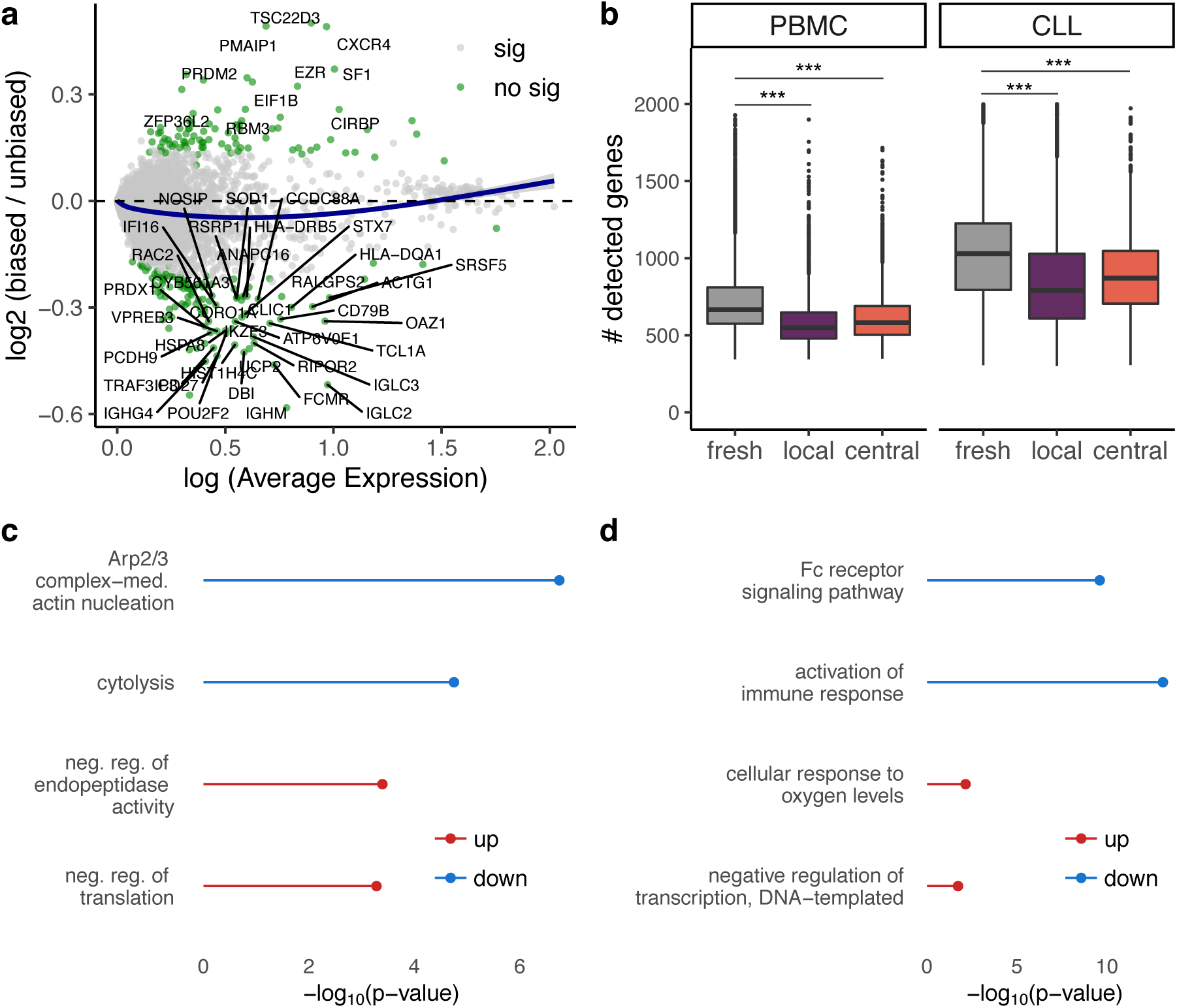
Cold-shock scRNA-seq signature. (**a**) M(log ratio)-A(mean average) plot showing the log_2_ fold-change between biased (>2h) and unbiased (<=2h) CLL cells as a function of the log average expression (*Scran* normalized expression values). Significant genes with an adjusted p-value <0.001, an absolute log2 fold-change >0.25 and a log (average expression) >0.5 are colored in green. (**b**) Distribution of the number of detected genes across processing types (fresh, local, central) in both PBMC and CLL cells. *p<0.05, **p<0.01, ***p<0.001. (**c**,**d**) Gene ontology enrichment analysis of the up- and down-regulated genes in the cold-shock signatures in PBMC (**c**) and CLL (**d**) cells.

**Supplementary Figure 7.**
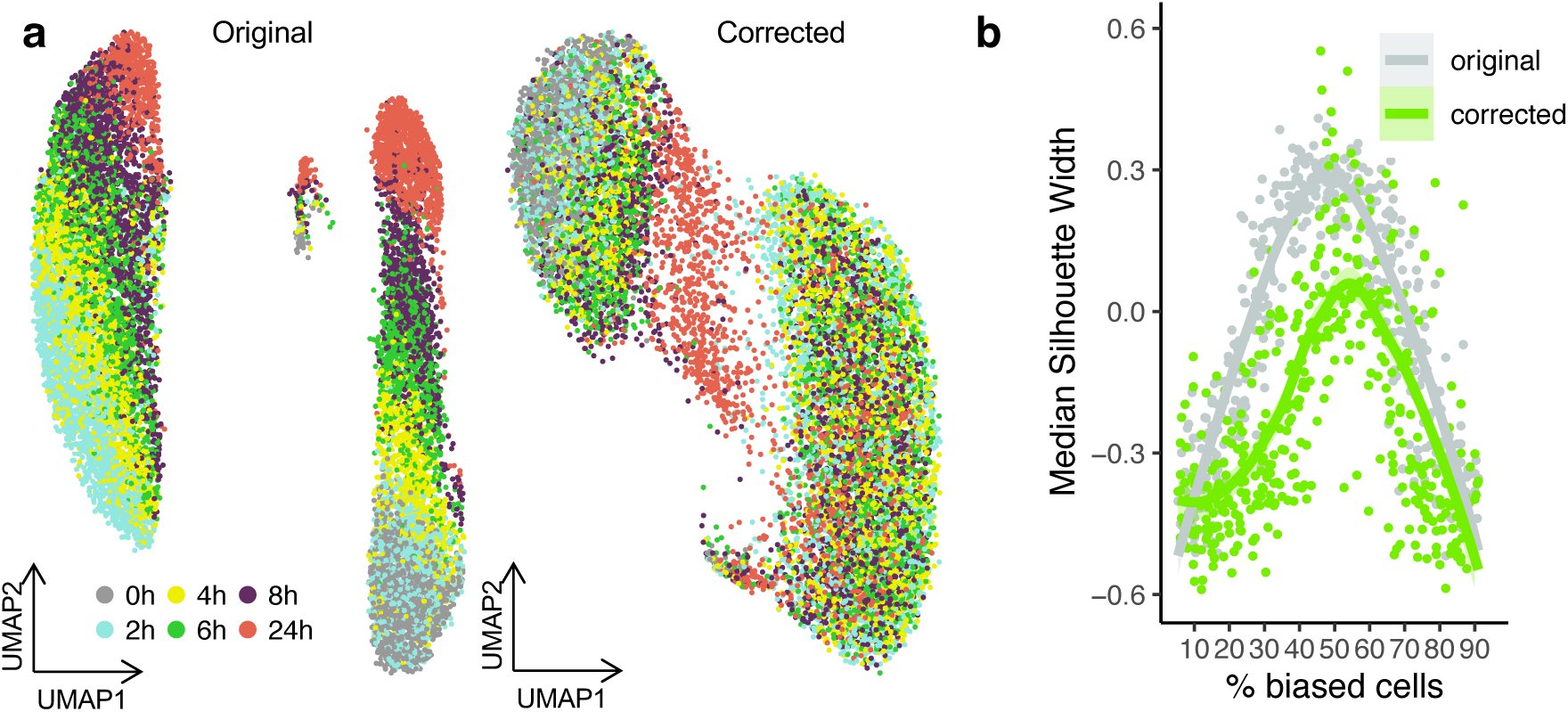
Cold-shock scRNA-seq signature correction. **(a)** tSNE embedding of CLL cells from two donors before (left) and after (right) regressing out the cold-shock score. **(b)** Assessment of the computational correction robustness by bootstrapping datasets with varying compositions of affected cells. Silhouette width is inversely proportional to the intermixing of the conditions.

**Supplementary Figure 8.**
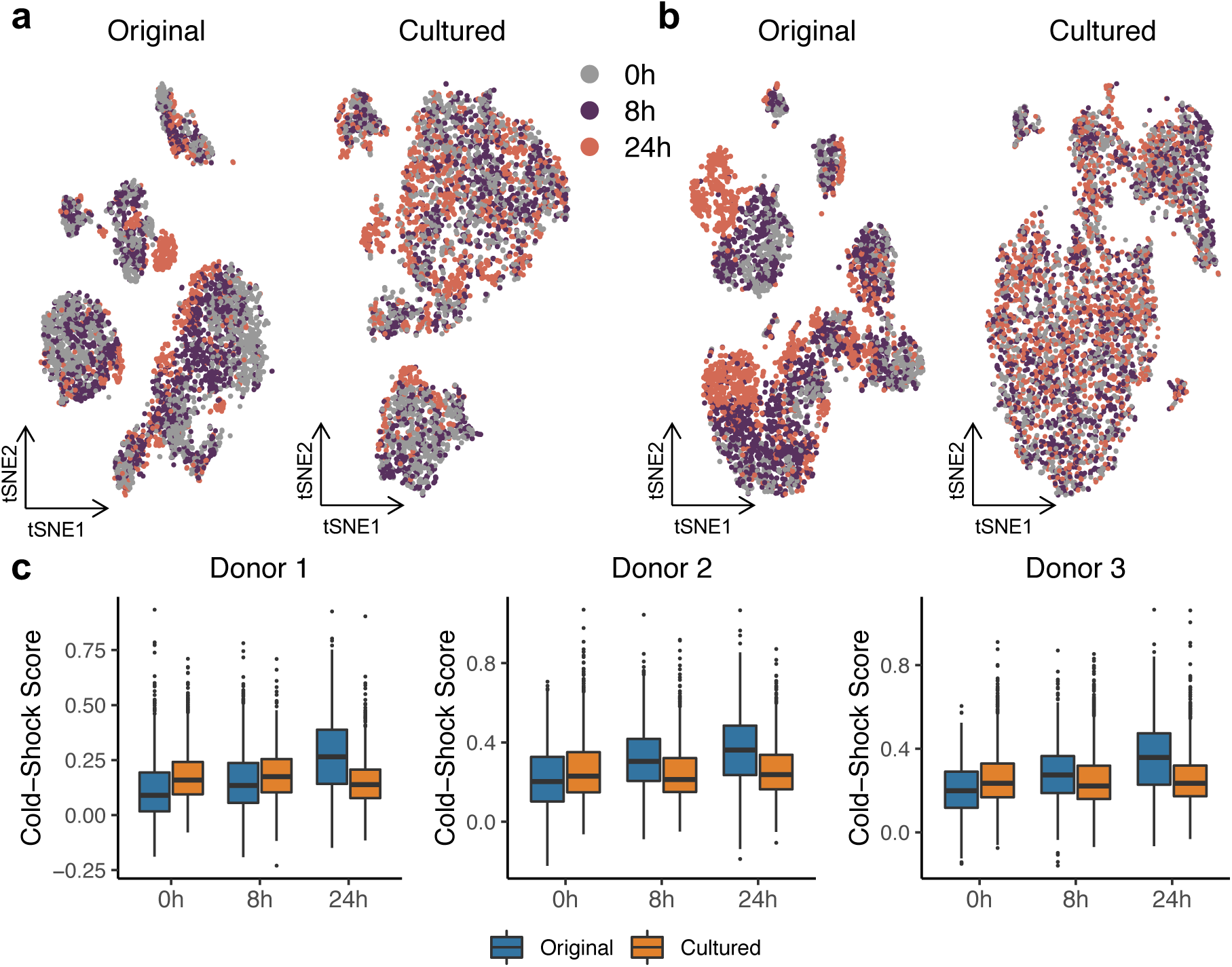
Culturing PBMC removed the sampling time-associated bias. (**a, b**) tSNEs showing the effect of culturing and CD3-activation of PBMC over two days on donors 2 (a) and 3 (b). (**c**) Cold-shock distribution across sampling times with (orange) or without (blue) culturing and activation of PBMC prior to processing.

**Supplementary Figure 9.**
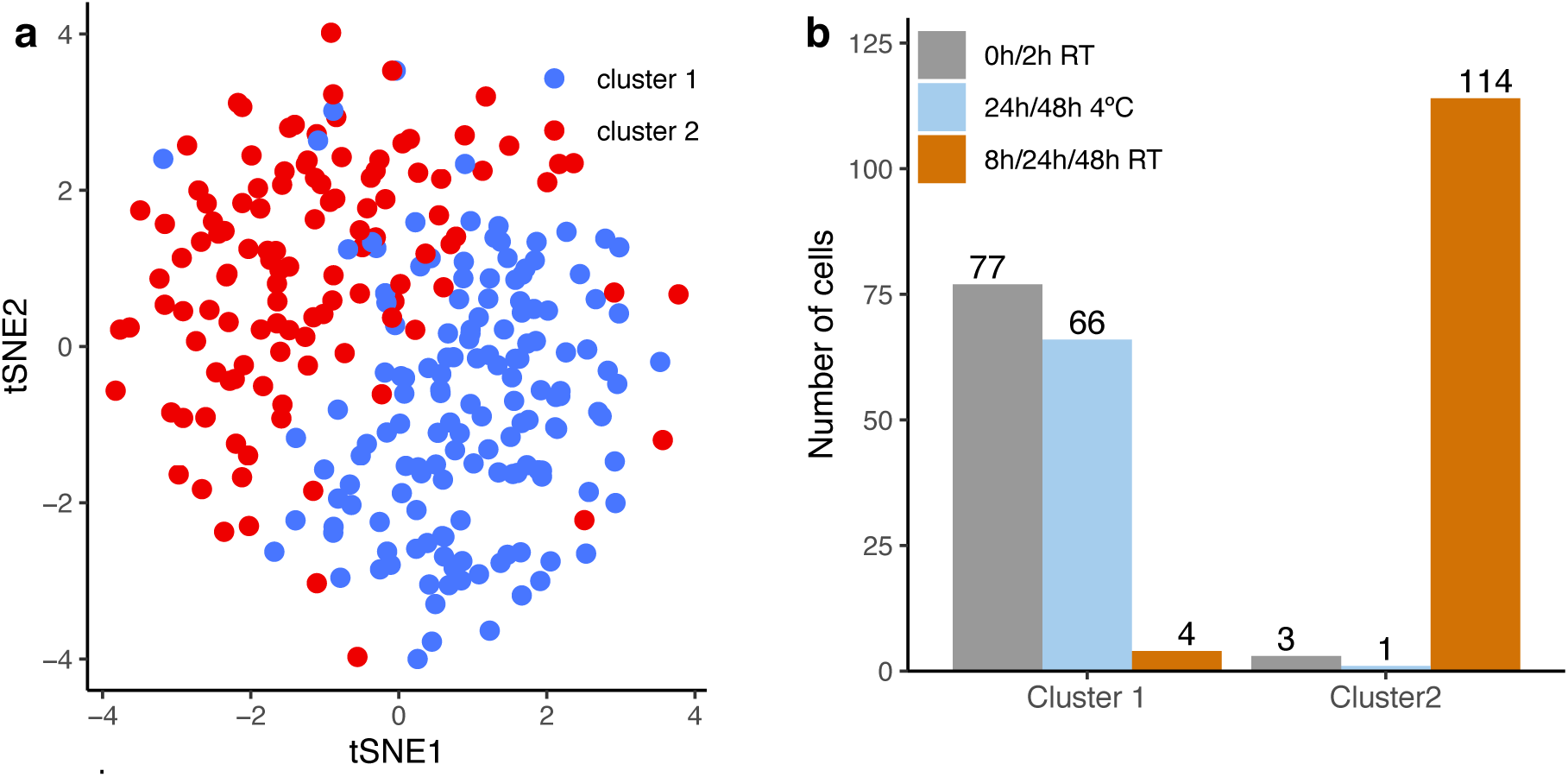
The impact of sampling time at 4°C on scRNA-seq profiles. (**a**) tSNE embedding of 265 CD3+ cells cryopreserved (i) immediately after blood extraction, (ii) storage at 4°C or (iii) at 21°C for 24/48 h prior to cryopreservation. Cells are colored by cluster label (SC3). (**b**) Distribution of processing conditions across cell clusters.

**Supplementary Figure 10.**
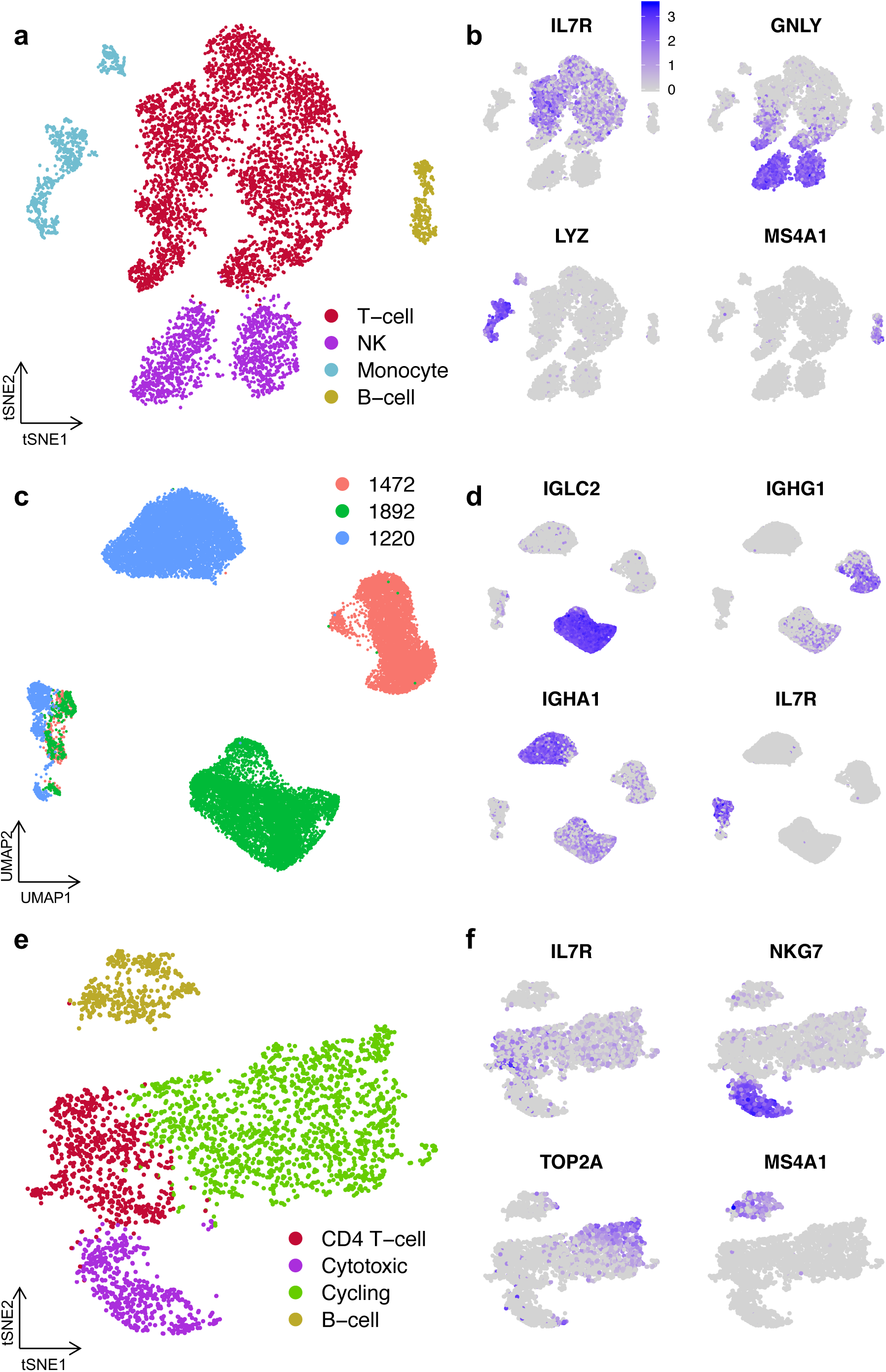
**(a**,**c**,**e)** tSNE embedding of (**a**) PBMC, (**c**) CLL and (**e**) cultured PBMC colored by cell annotation. (**b**,**d**,**f**) tSNEs showing the selected markers for each of the cell annotations.

## References

1. Peakman, T. C. & Elliott, P. The UK Biobank sample handling and storage validation studies. Int. J. Epidemiol. 37 Suppl 1, i2-6 (2008).

2. Elliott, P., Peakman, T. C. & UK Biobank. The UK Biobank sample handling and storage protocol for the collection, processing and archiving of human blood and urine. Int. J. Epidemiol. 37, 234–244 (2008).

3. Guillaumet-Adkins, A. et al. Single-cell transcriptome conservation in cryopreserved cells and tissues. Genome Biol. 18, 45 (2017).

4. Puente, X. S., Jares, P. & Campo, E. Chronic lymphocytic leukemia and mantle cell lymphoma: crossroads of genetic and microenvironment interactions. Blood 131, 2283–2296 (2018).

5. Baechler, E. C. et al. Expression levels for many genes in human peripheral blood cells are highly sensitive to ex vivo incubation. Genes Immun. 5, 347–353 (2004).

6. Zhu, X., Bührer, C. & Wellmann, S. Cold-inducible proteins CIRP and RBM3, a unique couple with activities far beyond the cold. Cell. Mol. Life Sci. 73, 3839–3859 (2016).

7. Spriggs, K. A., Bushell, M. & Willis, A. E. Translational Regulation of Gene Expression during Conditions of Cell Stress. Mol. Cell 40, 228–237 (2010).

8. Al-Fageeh, M. B. & Smales, C. M. Control and regulation of the cellular responses to cold shock: the responses in yeast and mammalian systems. Biochem. J. 397, 247–259 (2006).

9. Brink, S. C. van den et al. Single-cell sequencing reveals dissociation-induced gene expression in tissue subpopulations. Nat. Methods 14, 935–936 (2017).

10. Tirosh, I. et al. Dissecting the multicellular ecosystem of metastatic melanoma by single-cell RNA-seq. Science 352, 189–196 (2016).

11. Scialdone, A. et al. Computational assignment of cell-cycle stage from single-cell transcriptome data. Methods 85, 54–61 (2015).

12. Büttner, M., Miao, Z., Wolf, F. A., Teichmann, S. A. & Theis, F. J. A test metric for assessing single-cell RNA-seq batch correction. Nat. Methods 16, 43–49 (2019).

13. Julious, S. A. & Mullee, M. A. Confounding and Simpson’s paradox. BMJ 309, 1480–1481 (1994).

14. Solovieff, N., Cotsapas, C., Lee, P. H., Purcell, S. M. & Smoller, J. W. Pleiotropy in complex traits: challenges and strategies. Nat. Rev. Genet. 14, 483–495 (2013).

15. Madissoon, E. et al. Lung, spleen and oesophagus tissue remains stable for scRNAseq in cold preservation. bioRxiv 741405 (2019) doi:10.1101/741405.

16. Stoeckius, M. et al. Cell Hashing with barcoded antibodies enables multiplexing and doublet detection for single cell genomics. Genome Biol. 19, 224 (2018).

17. Stuart, T. et al. Comprehensive Integration of Single-Cell Data. Cell 177, 1888-1902.e21 (2019).

18. Luecken, M. D. & Theis, F. J. Current best practices in single-cell RNA-seq analysis: a tutorial. Mol. Syst. Biol. 15, e8746 (2019).

19. McGinnis, C. S., Murrow, L. M. & Gartner, Z. J. DoubletFinder: Doublet Detection in Single-Cell RNA Sequencing Data Using Artificial Nearest Neighbors. Cell Syst. 8, 329-337.e4 (2019).

20. L. Lun, A. T., Bach, K. & Marioni, J. C. Pooling across cells to normalize single-cell RNA sequencing data with many zero counts. Genome Biol. 17, 75 (2016).

21. Satija, R., Farrell, J. A., Gennert, D., Schier, A. F. & Regev, A. Spatial reconstruction of single-cell gene expression data. Nat. Biotechnol. 33, 495–502 (2015).

22. McCarthy, D. J., Campbell, K. R., Lun, A. T. L. & Wills, Q. F. Scater: pre-processing, quality control, normalization and visualization of single-cell RNA-seq data in R. Bioinformatics 33, 1179–1186 (2017).

23. Soneson, C. & Robinson, M. D. Bias, robustness and scalability in single-cell differential expression analysis. Nat. Methods 15, 255–261 (2018).

24. Vieth, B., Parekh, S., Ziegenhain, C., Enard, W. & Hellmann, I. A systematic evaluation of single cell RNA-seq analysis pipelines. Nat. Commun. 10, 1–11 (2019).

25. Falcon, S. & Gentleman, R. Using GOstats to test gene lists for GO term association. Bioinformatics 23, 257–258 (2007).

26. Picelli, S. et al. Smart-seq2 for sensitive full-length transcriptome profiling in single cells. Nat. Methods 10, 1096–1098 (2013).

27. Kiselev, V. Y. et al. SC3: consensus clustering of single-cell RNA-seq data. Nat. Methods 14, 483–486 (2017).

28. Kiselev, V. Y., Andrews, T. S. & Hemberg, M. Challenges in unsupervised clustering of single-cell RNA-seq data. Nat. Rev. Genet. 20, 273 (2019).

29. Kopp, W. & Vingron, M. An improved compound Poisson model for the number of motif hits in DNA sequences. Bioinforma. Oxf. Engl. 33, 3929–3937 (2017).

